# A new semi-automated, motility-based screening assay for discovery of compounds with activity against the juvenile stage of *Fasciola hepatica*

**DOI:** 10.64898/2026.06.22.733915

**Authors:** Alice Bernal, Diana S. Gliga, Giulia Colangeli, Matías Preza, Rossitza N. Irobalieva, Caroline F. Frey, Andrew Hemphill, Britta Lundström-Stadelmann, Natalie Wiedemar

**Author notes:** Sección Biología Celular, Facultad de Ciencias, Universidad de la República, Montevideo, Uruguay (Matías Preza).

## Abstract

*Fasciola hepatica* is a trematode parasite responsible for fasciolosis, a liver disease that affects humans and livestock worldwide. Together with other food-borne trematode infections, fasciolosis is considered a neglected tropical disease. Further, it imposes substantial agricultural losses due to infections in ruminants. No vaccine is currently available, and control heavily relies on drug treatment, especially with triclabendazole (TCBZ). However, the intensive use of TCBZ over the past four decades has led to increasing rates of treatment failures and the emergence of drug-resistant parasites. Therefore, the identification of new treatment options is an urgent priority.

The currently available toolset for drug screening, however, is limited. To address this need, we established a novel, semi-automated, standardized, and objective screening assay based on motility monitoring of newly excysted juveniles using microscopic live imaging. The assay was validated by testing a panel of ten compounds with known anthelmintic properties, amongst them TCBZ (IC_50_: 1.5 µM) and the new activator of the *F. hepatica* transient receptor potential melastatin (TRPM) ion channel, benzamidoquinazolinone (IC_50_: 1.05 µM). In addition to these two compounds with known activity against *F. hepatica*, three compounds were identified as particularly promising with a fast onset of action and IC_50_ values in the nanomolar range: the salicylanilides MMV665807 (IC_50_: 44 nM), niclosamide (IC_50_: 32 nM), and its ethanolamine salt, niclosamide ethanolamine (IC_50_: 9 nM). Complementary live/dead staining revealed that only TCBZ displayed parasiticidal activity, while the other compounds, although leading to parasite paralysis, did not lead to parasite death within 72 hours. Scanning electron microscopy of drug treated parasites did not reveal any significant damage at concentrations corresponding to the IC_50_s, but strong phenotypes were visible at 20 µM.

The presented motility assay provides a robust method for the discovery of novel anthelmintic compounds and facilitates the ongoing effort to combat fasciolosis.

**Author Summary:** *Fasciola hepatica*, the common liver fluke, is a parasitic platyhelminth that infects the liver and biliary ducts of humans and livestock, causing fasciolosis, a Neglected Tropical Disease as defined by the World Health Organization. Triclabendazole is the drug of choice to treat humans and animals. However, its intensive use has led to the emergence of drug resistance resulting in treatment failures worldwide. The identification of novel drugs is therefore urgent. Here, we present a semi-automated and objective method to assess the activity of compounds on one of the key life stages of the parasite: the newly excysted juveniles (NEJ). This stage is highly motile and motility assessment can be exploited to screen for bioactive compounds. Using time-lapse imaging, we quantified NEJ movement after drug exposure. From a panel of ten tested reference anthelmintics, two known fasciolicides (triclabendazole and benzamidoquinazolinone) and three additional compounds (MMV665807, niclosamide, and niclosamide ethanolamine) displayed particularly strong activity and were selected for further investigation.

This method represents a robust tool for drug screening and facilitates the discovery of new compounds against *F. hepatica*.

## Introduction

Fasciolosis is a parasitic disease affecting the liver. It is caused by the globally distributed trematode *Fasciola hepatica*, a member of the Fasciolidae family, also called the common liver fluke. Human fasciolosis is a recurrent issue in low-income countries, with poorer population groups and children particularly at risk due to limited access to safe water and the frequent consumption of raw aquatic plants [1]. The Peruvian and Bolivian highlands are the best-known high endemicity regions, several studies reported alarming infection rates in the past, such as in the Cajamarca province in Peru, with prevalences of 6.7% to 47.7% in 2-18 year old school children [2], or in the Los Andes province in the Northern Bolivian Altiplano with prevalences of up to 67% [3]. Consequently, fasciolosis was classified as a Neglected Tropical Disease by the World Health Organization [4]. Besides the human health burden, fasciolosis represents a major, global agricultural concern. Infections in livestock have significant economic impact due to decreased meat and milk production, reduced fertility, and liver condemnations at the abattoir. In Europe, for instance, fasciolosis is responsible for an estimated € 635 million of agricultural loss per year [5].

Mammalian infection occurs through ingestion of metacercariae that are commonly found on water plants in freshwater habitats harboring the intermediate host, lymnaeid snails. After ingestion, metacercariae excyst in the small intestine and juvenile flukes penetrate the intestinal wall and migrate to the liver. There, they traverse the hepatic parenchyma over a period of 5–6 weeks before entering the bile ducts, where they mature into adult flukes and begin sexual reproduction. Eggs excreted by the adults are released via the bile and feces into the environment. In freshwater, miracidia mature and hatch from the eggs, they swim, actively seek and infect lymnaeid snails. Within the snail host, the parasites undergo asexual reproduction and are eventually released back into the water as cercariae. These free-swimming forms encyst on submerged vegetation or other surfaces, transforming into metacercariae, ready to infect a new mammalian host and complete the life cycle [6]. The migration of juvenile flukes through the liver parenchyma is typically associated with acute abdominal and hepatic symptoms, including fever, hepatomegaly, abdominal pain and hepatitis. This initial phase, known as the hepatic stage, corresponds to the first clinical manifestation of fasciolosis [7]. Once the parasites reach and establish in the bile ducts, they provoke more chronic symptoms, primarily due to mechanical obstruction of the bile flow. This marks the second clinical stage, referred to as the biliary phase [7]. The infection is diagnosed through clinical examination, coproscopy, coproantigen testing and serology.

As no vaccine is currently available, control relies heavily on drug treatment. Triclabendazole (TCBZ), a member of the benzimidazoles, is the most frequently used drug against fasciolosis [8] as it is active against both the adult and the juvenile flukes. It is the only drug approved for use in humans and proposed to target β-tubulin similar as other benzimidazoles. Several other drugs are available for veterinary use [8,9] such as the benzimidazole albendazole (targeting β-tubulin), the sulphonamide clorsulon that inhibits phosphoglycerate kinase [10], the phenol nitroxynile, and the salicylanilides closantel and oxyclozanide which are all uncouplers of oxidative phosphorylation but might have additional targets and mode of actions [9]. However, the activity of these compounds is limited to the more mature stages of the parasite. The intensive use of TCBZ over the past four decades has resulted in increasing rates of treatment failures and the emergence of drug-resistant parasites. The first resistant *F. hepatica* were reported from Australia [11], followed by a still increasing number of reports about TCBZ resistance from multiple countries across Europe and South America [12]. These reports were predominantly from veterinary medicine, however, recently, TCBZ treatment failure has also been observed in human patients in Peru, Egypt, and Portugal [13–15]. For instance, in Peru, a concerning number of treatment failures was reported among children, with 10% remaining uncured despite undergoing multiple rounds of high-dose therapy [16]. Recent studies identified multiple genomic loci associated with resistance in British and Peruvian isolates [17,18], however, the underlying mechanisms of resistance remain poorly understood.

For these reasons, the discovery of new therapeutic options is crucial. A limited number of compound screens targeting *Fasciola* spp. have been reported, applying either target-based approaches or whole organism assays. The former requires a well-characterized drug target and an enzymatic assay to test potential drug candidates for *Fasciola*, such as the cysteine protease cathepsin L [19,20], the platyhelminth specific thioredoxin glutathione reductase [21], or UDP-galactose 4’-epimerase [22]. Whole organism based assays assess the effect of compounds on whole parasites and have in the past been conducted on metacercariae [23], newly excysted juveniles (NEJ, [24]), juveniles, adults and eggs [25,26]. Read-outs of these assays have typically been alterations of morphology, inhibition of motility or inhibition of egg embryonation as assessed by visual and microscopic inspection and manual scoring. While these assays provide valuable information about the activity of individual compounds on different parasite life forms, their objectivity and scalability are limited due to the manual readouts. Keiser et al. used microcalorimetry to measure the heat-flow of juvenile and individual adult *F. hepatica* exposed to selected compounds [27], which provided detailed and objective parasite viability time-course data, and Zheng et al. recently developed a larval screening method with miracidia using the infrared detection system WMicroTracker ONE providing automatic and scalable read-outs [28]. As the juvenile and adult *F. hepatica* parasite forms are responsible for pathology in the mammalian host, we aimed to develop a scalable screening method with one of these stages. Adult-stage screening presents significant limitations: parasites are big (up to 3 cm long), not available in large numbers and assay setup is complex and labor-intensive, which considerably reduces throughput and does not allow for larger-scale compound testing. NEJ parasites instead can be obtained at higher numbers by excystation of metacercariae and used in scalable setups such as on 384-well plates. Therefore, we focused on the NEJ stage of *F. hepatica*. A key characteristic of NEJ is their motility that allows migration from the intestinal lumen via peritoneum to and through the liver tissue [29]. Consequently, motility can be used to identify bioactive compounds that either paralyze or kill the parasites. We adapted an imaging-based motility assay applied previously on *Echinococcus multilocularis* and *E. granulosus* protoscoleces [30,31], as well as on *Taenia crassiceps* cysticerci [32], for use in *F. hepatica*. We evaluated the assay on a panel of ten compounds with reported activities against platyhelminths. Amongst them, we tested broad-spectrum benzimidazoles and praziquantel, the latter is known to be inactive in *F. hepatica* due to a point mutation in the drug binding site of its target, a transient receptor potential melastatin ion channel (TRPM_PZQ_ [33]), and served as a negative control. We further included benzamidoquinazolinone (BZQ, [34]), a recently discovered molecule which targets an ubiquitously conserved residue within TRPM_PZQ_ enabling activity of this compound against *F. hepatica*. Also, we included drug repurposing candidates such as niclosamide and selected compounds from the Medicines for Malaria Venture Pathogen Box that have demonstrated promising *in vitro* anti-helminthic activity [35–38] and TCBZ as positive control. Initially, we tested these compounds at high concentrations and compared the motility data with dual live-dead staining. We next assessed the most promising compounds by determining their 50% inhibitory concentrations (IC_50_) and visualized morphological changes induced by drug treatment using scanning electron microscopy (SEM). With this method, we present a first objective, semi-automated drug screening assay for *F. hepatica* NEJ, that will facilitate the identification of novel fasciolicidal compounds in the future and we identified three molecules with bioactivity on NEJ.

## Materials and methods

If not stated otherwise, all plastic ware was purchased from Sarstedt (Sevelen, Switzerland).

### Source of metacercariae

Metacercariae were either purchased from Ridgeway Research Limited, St Briavels, United Kingdom (TCBZ-sensitive Aberystwyth and Italian strains) or produced in house. Metacercariae were stored in sterile, distilled water in 50 mL tubes at 4°C, the water was replaced every two weeks.

### In-house production of metacercariae

Adult *F. hepatica* parasites were recovered from naturally infected bovine livers obtained at a local abattoir (Bell, Oensingen, Switzerland). Parasites were incubated overnight in RPMI 1640 (LabForce AG Muttenz, Switzerland) with 10% fetal calf serum (FCS, Seraglob, Bioswisstec, Schaffhausen, Switzerland) and 1× Antibiotic-Antimycotic solution (100×; Thermo Fisher Scientific, Waltham, MA, USA), at 37°C, 5% CO_2_ and the excreted eggs were collected and stored in distilled water at 4°C. For maturation of the miracidia, the eggs were incubated in the dark at 27°C. After 14-16 days, hatching of miracidia was induced through exposure to natural light and 30 µL aliquots of the miracidia-solution transferred to the wells of a 48-well plate. The wells were inspected under an inverted microscope for presence of two miracidia. 500 µL tap water and one in-house bread, uninfected *Galba truncatula* snail (∼4-5 mm shell length) were subsequently added to positive wells. Snails were left for 2-3 hours in the wells to allow infection to take place (room temperature and day light exposure). Infected snails were then transferred to and cultured in glass containers with 500-800 mL tap water and the common water moss *Fontinalis antipyretica* (*in vitro* culture purchased from Swiss Aqua Shop, Lausanne, Switzerland or from Garnelenmarkt, Bern, Switzerland) at room temperature, illuminated 12 hours / day with a LED Aquarium Lighting (WEAVERBIRD Gemini). The containers were cleaned three times a week and the snails fed with spirulina chips (Aqua Terra-Discount, Turgi, Switzerland) and plant pellets (25% nettle, 25% dandelion, 25% ribwort plantain, 25% spinach from Aqua Terra-Discount, Turgi, Switzerland). After 35 days, snails were transferred to smaller glass dishes containing 50 mL of tap water and lined with cellophane foil where they were allowed to shed cercariae naturally. Metacercariae were collected three times a week.

### Excystation

To prevent metacercariae and NEJ from sticking to surfaces, all plasticware and pipet tips were coated with FCS and rinsed with PBS before use. Metacercariae were excysted with a protocol adapted from Smith & Clegg, 1981 [39]. Briefly, they were first incubated for one hour at 39°C in the dark in 10 mL activation solution consisting of 20 mM sodium hydrosulfite (Sigma-Aldrich, CHEMIE, GmbH, Germany) prepared with cold CO_2_ enriched water and warmed up to 39°C directly before use. Subsequently, metacercariae were washed twice with 39°C PBS and eluted in 10 mL pre-warmed excystation medium consisting of Hank’s Balanced Salt Solution (Sigma-Aldrich, CHEMIE, GmbH, Germany), 30 mM HEPES, and 10% sheep bile (obtained by sterile puncture of gall bladders obtained from a local abattoir; and stored frozen at −20°C). The excystation media containing the metacercariae was transferred to petri dishes and incubated for up to three hours at 39°C, 5% CO_2_ in a humidified incubator in the dark. NEJ were collected after one, two and three hours of incubation and transferred to pre-warmed NEJ serum-free media (NEJ-SF-media) consisting of RPMI 1640 supplemented with 1x Antibiotic-Antimycotic solution and 1x Glutamax (Gibco, Thermo Fisher Scientific). Upon completion of collection, NEJ were washed 2x with NEJ-SF-media to remove residual bile from the excystation medium.

### Motility Assay: Plate setup and imaging

Directly after excystation, NEJ were suspended in NEJ-SF-media and distributed on a 384-well plate (Corning 3701 supplied by Fisher Scientific, Reinach, Switzerland) in a volume of 60 µL per well. The plate was incubated for 24 hours in a humidified incubator at 37°C, 5% CO_2_ and parasite motility quantified by live imaging (see below “Quantification of the motility by live imaging”). To investigate the effect of different imaging intervals (5, 10, 20, 30 and 50 s) on motility read-outs, wells with 5-7 parasites were imaged for 4 min. Images were taken every 5 seconds and the different intervals compared (see below “Quantification of motility by live imaging”).

To study the effect of the parasite numbers on motility read-outs, wells with 1, 2, 3, 4, 5, 7 and 10 parasites were imaged with the setup 4 pictures, 10 seconds apart.

### DMSO and triclabendazole assays

After excystation, NEJ were added to a 384-well plate with five NEJ in 30 µL NEJ-SF-media per well. Medium-drug solutions were prepared with TCBZ or DMSO at 2x the final concentrations. 30 µL of the media-drug solution were added to the wells with the parasites resulting in final concentrations of 20, 10, 7, 5, 3.5, 2.5 and 1.25 µM. Every concentration was tested in duplicate on each plate. As controls, we included 0.4% DMSO (corresponding to the highest DMSO concentration used in the assay) and medium only. After addition of the drugs, the plates were incubated in a humidified incubator at 37°C, 5% CO_2_. After 6, 12, 24, 48 and 72 hours, the parasite motility was quantified by live imaging as described below.

### Assessment of anthelmintic compounds

Five NEJ suspended in 30 µL NEJ-SF-media were added to the wells of a 384-well plate. The following compounds were tested: albendazole (Merck, Buchs, Switzerland), mebendazole (Lucerna-Chem, Lucerne, Switzerland), fenbendazole (Merck, Buchs, Switzerland), praziquantel (Novartis Pharma Schweiz AG, Rotkreuz, Switzerland), niclosamide (Sigma-Aldrich, CHEMIE, GmbH, Germany), niclosamide ethanolamine (NEN; from Biotech, USA), MMV665807 (Evotec, USA), MMV689480 / buparvaquone (Evotec, USA), BZQ (provided by Daniel Sprague & Jonathan Marchant, Department of Cell Biology, Neurobiology, and Anatomy, Medical College of Wisconsin, Milwaukee, WI, USA [34]), and TCBZ (Merck, Buchs, Switzerland). Compounds were eluted at 10 mM in ≥99% dimethylsulfoxide (DMSO), TCBZ was prepared at 5 mM in ≥99% DMSO. Drug-media solutions for each of the ten compounds were prepared at 20 µM and 80 µM in NEJ-SF-media. 30 µL of the drug-media solutions were added to the wells with the NEJ resulting in final concentrations of 10 µM and 40 µM and a total volume of 60 µL per well. Drugs were tested in duplicate on each plate. 0.4% DMSO was used as control, as well as NEJ-SF-media. After addition of the drugs, the plate was incubated in a humidified incubator at 37°C, 5% CO_2_. After 24, 48 and 72 hours, parasite motility was quantified by live imaging as described below.

### Dose response curves

30 µL of NEJ-SF-media with 5 parasites were added to the wells of a 384-well plate. Drug solution series of NEN, MMV665807, niclosamide, BZQ and TCBZ were prepared in NEJ-SF-media and 30 µL added to the wells with the parasites. Drugs were tested in duplicate on each plate. 0.4% DMSO was used as control, as well as NEJ-SF-media. After addition of the drugs, the plate was incubated in a humidified incubator at 37°C, 5% CO_2_. After 72 hours, parasite motility was quantified by live imaging as described below. Dose-response curves were fitted with a 4-parameter log-logistic dose-response model with Gaussian default for continuous responses, nonlinear least squares in Rstudio v 4.4.1.

### Quantification of motility by live imaging

A series of images was taken of each well at defined time intervals (final setup: 4 pictures, 10 seconds apart) using a Nikon Ti2 inverted microscope, with a Hamamatsu Orca-fusion camera, with the software NIS Elements AR, version 6.20.02, 64-bit. Imaging took place in a heated chamber at 37°C, 5% CO_2_.

ImageJ was used to subtract pixel intensities of each frame from the pixel intensities of the previous frame for each image series. The size of the area that differed between the images was then quantified, resulting in the raw motility in Δ pixels. The mean of the raw motilities of each image series was divided by the number of parasites per well resulting in the “Average Motility per NEJ” (in Δ pixels; RStudio version 4.4.1). Image acquisition and analysis are schematically depicted in Fig S1. To quantify the effect of the drugs on the parasites, the Average Motility per NEJ of drug treated parasites was divided by the Average Motility per NEJ of the medium only control resulting in percentage compared to the control.

### Live-dead staining

To differentiate between live and dead parasites, we used double staining with fluorescein diacetate (FDA) and propidium iodide (PI) (both purchased from Thermo Fisher Scientific AG, Basel, Switzerland) at the assay endpoint after 72 hours of incubation with the drugs (see above “Assessment of anthelmintic compounds”). FDA diffuses through membranes and is hydrolysed intracellularly to free fluorescein which cannot leave intact membranes due to its polarity, thereby it selectively stains living cells. PI instead, crosses damaged membranes and binds to DNA, thereby labelling dead cells. We prepared a 2× staining solution containing 8 µg/mL of PI and 1 µg/mL of FDA in PBS. The parasites in the assay plate were washed three times with 37°C PBS. After the last wash, the staining solution was added to a final concentration of 4 µg/mL PI and 0.5 µg/mL FDA in a total volume of 60 µL per well. The plate was incubated at 37°C, 5% CO_2_ for 20 minutes in the dark and imaged on a Nikon Ti2 Microscope using NIS Elements. PI staining was detected at 550 nm excitation and 561 nm emission, and FDA staining at 475 nm excitation and 488 nm emission.

### Scanning Electron Microscopy

Drug-NEJ-SF-media solutions were prepared with NEN, niclosamide, NEN, MMV665807, and BZQ concentrations corresponding to their IC_50_s (0.69 µM, 0.01 µM, 0.03 µM and 0.04 µM respectively) and at 20 µM. Approximately 20 NEJ were added to each solution and incubated for 72 hours at 37°C. The NEJ-SF-media containing the drugs was subsequently removed and the parasites washed 2× with PBS in a 1.5 mL screw-cap tube. Parasites were centrifuged at 500 x g for 2 min, the supernatant was removed, and 1 mL 100 mM sodium cacodylate buffer, pH 7.3 was carefully added, without resuspending the parasites. All subsequent fixation, washing and dehydration steps were done without resuspension of parasites. Primary fixation was carried out with 2.5 % glutaraldehyde in cacodylate buffer overnight at 4°C. After 2 washes in cacodylate buffer, specimens underwent post-fixation in 2% osmium tetroxide in cacodylate buffer for 2 hours at 24°C, followed by one wash in water and stepwise dehydration in a graded series of ethanol (30-50-70-90 and 3×100%), 2-3 minutes each. Specimens were then washed twice in hexamethyl-disilazane (HMDS; Merck), and after removal of the liquid, they were air-dried at 24°C overnight. The very tips of the Eppendorf tubes were carefully removed with a scalpel, leaving a small cup, which was then placed onto SEM stubs. Specimens were sputter-coated with gold and were imaged on an Apero-2S SEM operated at 2kV with a T2 detector.

### Statistical tests

All analysis were performed using R Statistical Software and RStudio (v4.4.1; R Core team 2024)

## Results

### Assay optimization for quantification of *F. hepatica* NEJ movement

Live *F. hepatica* NEJ can move in various ways: they contract in length while staying on site, they show forward locomotion, and sometimes they reside still while only internal gut movements are visible (supplementary Video S1). To consider all types of movement, we quantified parasite motility by analysing changes of pixel intensities between frames in a time series of microscopic live images. We initially optimized several parameters for quantification of *F. hepatica* NEJ movement. We imaged parasites every 5 seconds and compared different time intervals (5, 10, 20, 30, or 50 seconds) to evaluate, if motility values increase with longer intervals (Fig 1A, B). Average Motility per NEJ varied across the wells and ranged from 877 ± 486 to 1981 ± 426 Δpixels. A one-way repeated-measures ANOVA across the Average Motility per NEJ (in Δ pixels) revealed no significant difference between frame intervals (*p*-value = 0.34; Fig 1A), demonstrating that longer time intervals (20, 30, and 50 seconds) did not result in higher motility scores compared to shorter intervals (5 and 10 seconds). Thus both, the 5- and 10-second intervals were sufficient for quantification of parasite motility. However, extending the interval to 10 seconds increases the likelihood of detecting movement in parasites with reduced motility (Fig 1B). Based on these findings, we selected a 10-second imaging interval for subsequent experiments, it allows a robust measurement and at the same time keeps the imaging duration manageable even for large numbers of tested wells and compounds.

**Fig 1.**
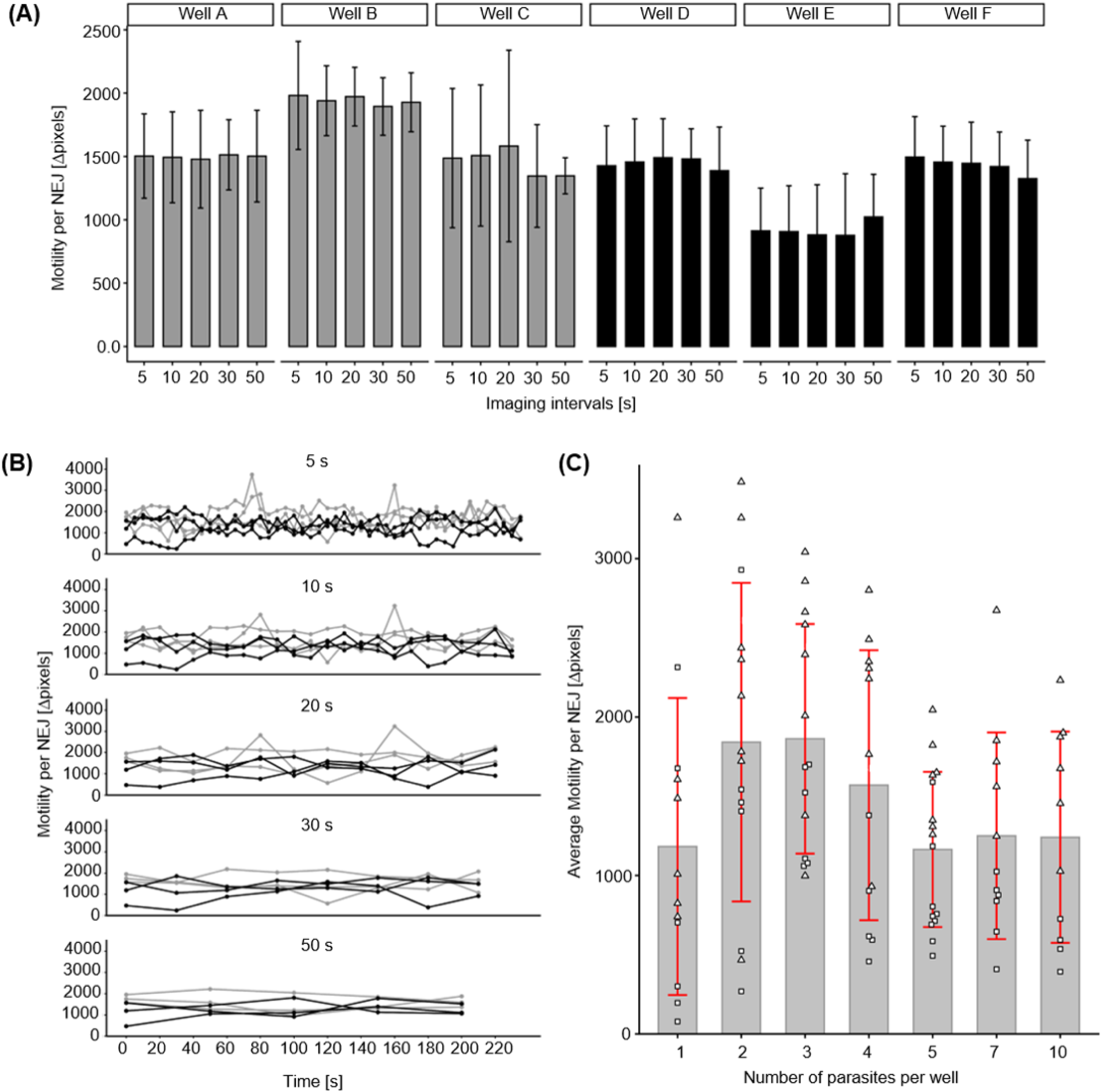
Effect of imaging interval duration and parasite number per well on quantification of *F. hepatica* NEJ movement. (A) Mean Motility per NEJ ± standard deviations are shown for individual wells as calculated with frame intervals of 5, 10, 20, 30, and 50 s. Data shown was derived from two biological replicates (parasites from two independent excystations) shown as black (replicate 1: well A, B, C) and grey (replicate 2: well D, E, F) bars. Every well contained between five and seven parasites and was imaged for 4 minutes. (B) Motility per NEJ over time for wells shown in (A). Individual replicates (wells) of the two independent experiments are represented in black and grey. (C) Average of 4 images taken 10 seconds apart (final setup) from wells containing 1, 2, 3, 4, 5, 7, and 10 parasites. Data is derived from two independent experiments (two excystations), individual replicates (wells) are shown as rectangle (experiment 1) and triangle (experiment 2) symbols. In each experiment at least 6 wells were imaged per condition (technical replicates: n ≥ 6). Bars represent the mean Motilities per NEJ across all replicates and error bars represent standard deviations.

To determine the minimum number of parasites required for consistent read-outs, we compared the Average Motility per NEJ across wells containing 1, 2, 3, 4, 5, 7, or 10 parasites (Fig 1C). For each condition, at least twelve wells were analysed (derived from two biological replicates). The mean Average Motility per NEJ (in Δ pixels) were 1’183 ± 937 for wells with 1 NEJ, 1’841 ± 1’005 for wells with 2 NEJ, 1’863 ± 725 for wells with 3 NEJ, 1’569 ±852 for wells with 4 NEJ, 1’164 ± 489 for wells with 5 NEJ, 1’250 ± 652 for wells with 7 NEJ, and 1’241 ± 666 for wells with 10 NEJ. Wells with only one or two parasites showed considerable variation, from three parasites onwards, standard deviations started to become smaller with more consistent clustering of motility values. Therefore, we established a minimum of five parasites per well as a requirement for robust motility scoring. The full process of image acquisition and analysis is depicted in Fig S1, for visualization of parasite movement and image subtraction see supplementary Video S1.

### Effect of DMSO and incubation time on parasite motility

Next, we assessed whether incubation in NEJ-SF-media alone or in NEJ-SF-media with DMSO had a (negative) impact on parasite movement over time. NEJ were incubated in NEJ-SF-media with or without 0.4% DMSO, which is the highest DMSO concentration we planned to use in the assay, parasite motility was quantified after 24, 48 and 72 hours (Fig 2A). The parasite movement increased significantly over time with an Average Motility per NEJ (in Δpixels) of 1’073 ± 493 after 24 hours, 1’357 ± 522 after 48 hours, and 1’633 ± 681 after 72 hours for parasites incubated in NEJ-SF-media (*p*-value NEJ-SF-media 24 h vs 72 h = 0.0003). A similar increase of motility was observed in parasites incubated in NEJ-SF-media with 0.4% DMSO, with an Average Motility per NEJ (in Δpixels)of 892 ± 328 after 24 hours, 1’271 ± 593 after 48 hours, and 1’348 ± 480 after 72 hours (*p*-value DMSO 24 h vs 72 h = 0.0257). Parasites incubated with and without DMSO did not show a significant difference if Average Motilities per NEJ were compared at the same time points.

**Fig 2.**
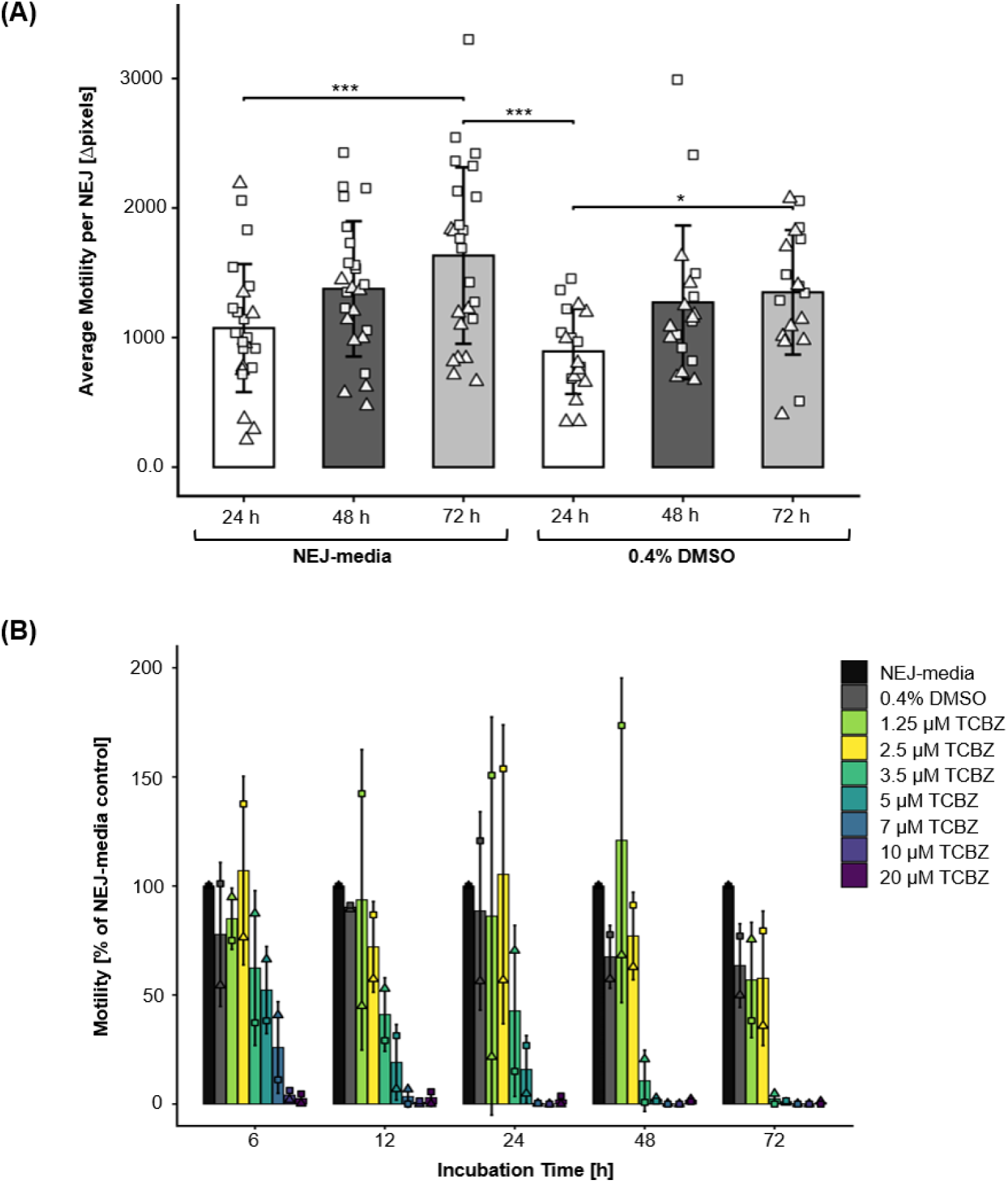
*F. hepatica* NEJ motility after incubation in NEJ-SF-media, DMSO and TCBZ. (A) Average Motilities per NEJ were determined after 24, 48, and 72 hours of incubation in NEJ-SF-media ± 0.4% DMSO. The dataset consists of two biological replicates (shown as rectangle and triangle symbols) derived from two independent excystations, each with eight technical replicates (eight wells) per condition. Bars display the mean and error bars the standard deviations across all replicates. Asterisks represent significant differences as calculated with an ANOVA followed by a Bonferroni-adjusted Tukey’s multiple-comparison test, *** = *p*-value ≤ 0.001, ** = *p*-value ≤ 0.01, * = *p*-value ≤ 0.05. (B) Average Motilities per NEJ normalized to the media-only control after incubation with different TCBZ concentrations for 6, 12, 24, 48, and 72 hours. Rectangles and triangles show the motility of two individual biological replicates (independent assays with juveniles from different excystations) each one consisting of two technical replicates (two wells). Bars represent the mean and error bars the standard deviations across biological replicates.

Next, we used TCBZ, to investigate dose-activity effects and to assess the optimal incubation time for detecting drug-induced effects (Fig 2B). TCBZ showed a relatively fast onset of action for higher concentrations with only residual motility remaining at the shortest exposure of 6 hours (3 ± 3% at 20 µM; 4 ± 3% at 10 µM as compared to the media-only control). At 12–24 hours ∼20% motility persisted at 5 µM (19 ± 17%, after 12h and 16 ± 16%, after 24h). After 48 hours, residual motility persisted at 3.5 µM (11 ± 14%). In contrast, after 72 hours, motility was completely abolished at 3.5 µM or higher, indicating that a 72-hour incubation time is necessary to increase the assay sensitivity and to reveal the full effect of the drug.

### Validation of the motility assay with a panel of anthelmintic compounds

We applied the *F. hepatica* NEJ motility assay on a panel of drugs and compounds with known anthelmintic properties to evaluate the performance of the method and to investigate the fasciolicidal potential of these compounds (Fig 3). Parasites exposed to the benzimidazoles albendazole, mebendazole and fenbendazole were still motile after 72 hours of incubation, even at the highest drug concentration of 40 µM with mean motility values of 83 ± 30% for albendazole, 74 ± 51% for mebendazole, and 56 ± 43% for fenbendazole. Similar results were observed for praziquantel with a residual motility of 62 ± 24% as compared to the no-drug control. These values did not significantly differ from the DMSO control that had a mean motility of 99.7 ± 36% after 72 hours (*p*-values *=* 0.99, 0.985, 0.452 and 0.698, respectively for albendazole, mebendazole, fenbendazole and praziquantel).

**Fig 3.**
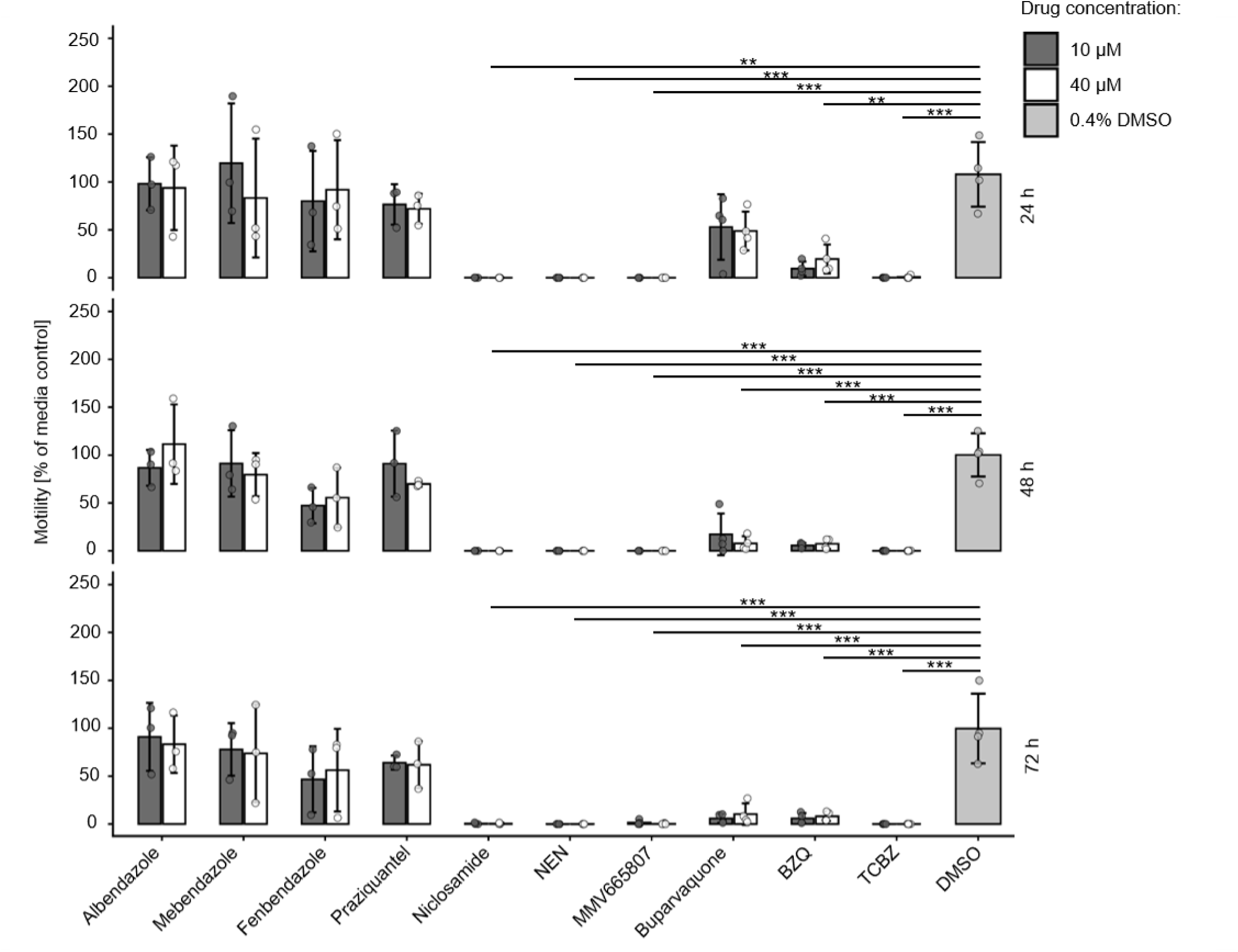
Effect of known anthelmintics on NEJ motility. Parasite motility after 24, 48, and 72 hours of incubation with albendazole, mebendazole, fenbendazole, praziquantel, niclosamide, NEN, MMV665807, buparvaquone, BZQ and TCBZ, each applied at 10 and 40 µM. Circles show the Average Motility per NEJ of individual biological replicates (independent assays, n = 3-4) each one consisting of two technical replicates (two wells), expressed as percentage of the media-only control. Bars represent the mean and error bars the standard deviations across biological replicates. Asterisks represent significant differences between the DMSO and 10 µM of the test compounds at 72 hours of incubation as calculated with an ANOVA followed by a Bonferroni-adjusted Tukey’s multiple-comparison test. Significance levels are displayed as *** = *p*-value ≤ 0.001, ** = *p*-value ≤ 0.01, and * = *p*-value ≤ 0.05.

In contrast, buparvaquone reduced the motility to 53 ± 34% at 10 µM, and to 49 ± 20% at 40 µM, after 24 hours. Motility further declined to 17 ± 22% at 10 µM, and 8 ± 7% at 40 µM after 48 hours and to 6 ± 5% at 10 µM, and 10 ± 11% at 40 µM after 72 hours, which significantly differed from the DMSO control (10 µM buparvaquone 72 hours vs DMSO *p*-value = 9.26e-6, 40 µM buparvaquone 72 hours vs DMSO *p*-value = 2.81e-5). This suggests a delayed onset of drug action, however, as values did not reach zero, the inhibition of NEJ motility by buparvaquone was not complete. Treatment with BZQ decreased NEJ motility to 9.5 ± 7%, and 19 ± 15%, at 10 and 40 µM after 24 hours treatment, to 6 ± 2%, and 7 ± 5%, after 48 hours, and to 6 ± 5%, and 8 ± 5%, after 72 hours of incubation significantly differing from the DMSO control (10 µM BZQ 72 hours vs DMSO *p*-value = 9.43e-6 and 40 µM BZQ 72 hours vs DMSO *p*-value *=* 1.61e-5) demonstrating that, even if motility was not completely abolished, BZQ strongly inhibited NEJ movement.

Niclosamide, NEN, MMV665807 and TCBZ completely abolished motility within 24 hours at 10 and at 40 µM, with motility values of 0% for all tested concentrations and timepoints, thus the results for all four compounds differed significantly from the DMSO control (after 72 hours - 10 µM niclosamide vs DMSO: *p*-value = 1.54e-5, 40 µM niclosamide vs DMSO: *p*-value = 1.52e-5, 10 µM NEN vs DMSO: *p*-value = 2.19e-6, 40 µM NEN vs DMSO: *p*-value = 2.19e-6, 10 µM MMV665807 vs DMSO: *p*-value = 3.16e-6, 40 µM MMV665807 vs DMSO: *p*-value = 2.33e-6, 10 µM TCBZ vs DMSO: *p*-value = 2.22e-6, 40 µM TCBZ-DMSO: *p*-value = 2.23e-6).

### Viability of drug-treated NEJs

To test whether drug-induced loss of motility reflects parasite death, we conducted live/dead double-staining with FDA and PI in one of our assays directly after the final imaging read-out following 72 hours of drug exposure (Fig 4, Supplementary Data Fig S2). TCBZ was the only drug that caused death of all parasites at 40 µM, and death of 6 out of 11 parasites at 10 µM as indicated by positive PI staining and absence of FDI staining. In contrast, for all other tested compounds, the majority of parasites remained viable after 72 hours of incubation even at 40 µM, as evidenced by positive FDA staining and absence of PI staining. This shows that niclosamide, NEN, MMV665807, and BZQ did not induce substantial parasite death after 72 hours of incubation despite significant impairment of parasite motility.

**Fig 4.**
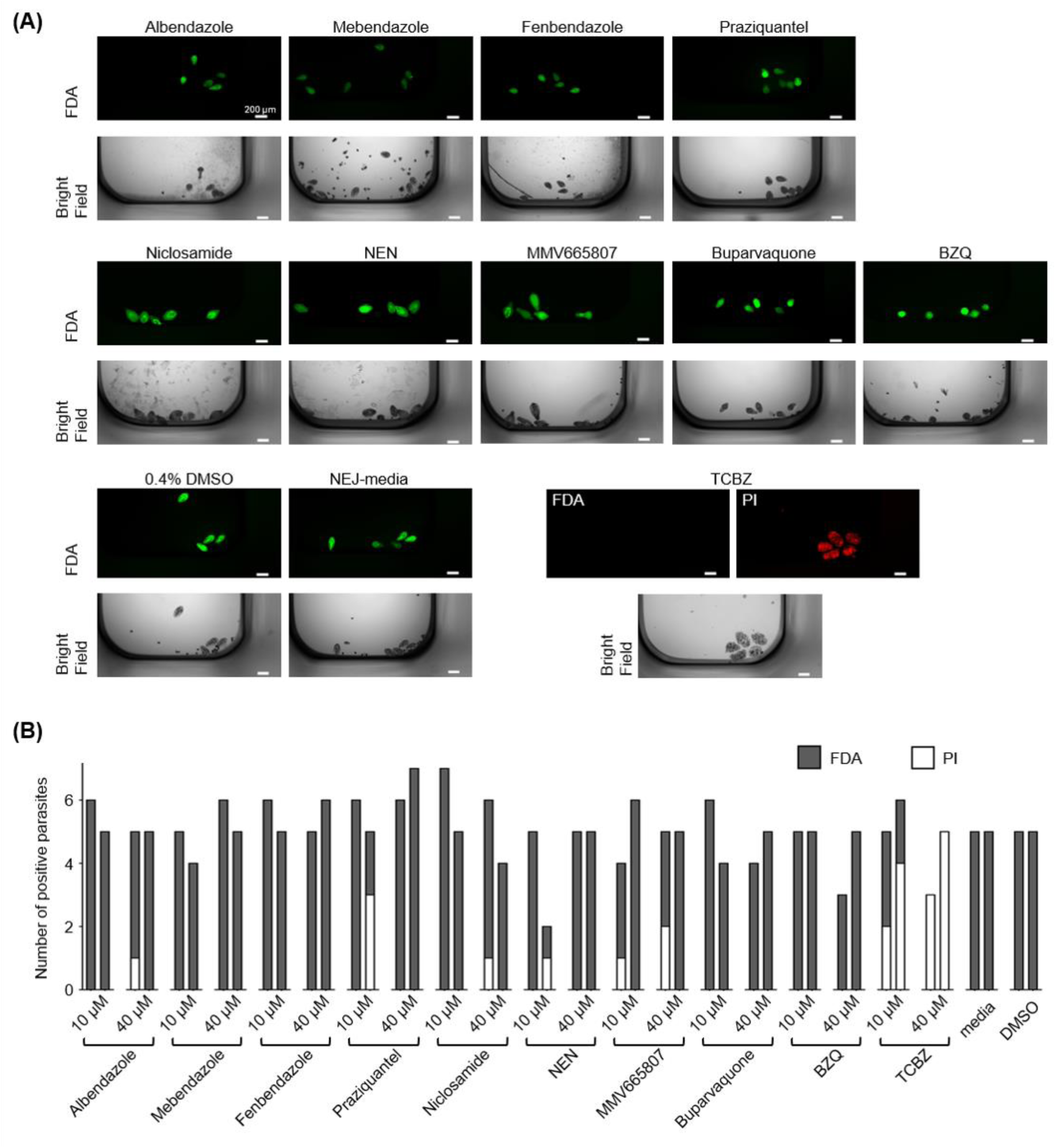
NEJ viability after 72 hours of drug exposure. (A) Representative images of FDA (green) and PI (red) double-stained parasites after incubation at 40 µM of the respective drug (all replicates are shown in Fig S2). Scale bars correspond to 200 µm. (B) Number of parasites stained positive with FDA or PI is shown with each bar representing an individual technical replicate (two wells per condition; source data in Fig S2).

### Potency of the active compounds and drug-induced morphological parasite alterations

We next assessed the dose-activity relationships of niclosamide, NEN, MMV665807, BZQ, and TCBZ on parasite movement after 72 hours of drug exposure. The most active compound was NEN with an IC_50_ of 9 nM followed by niclosamide with an IC_50_ of 32 nM. MMV665807 had an IC_50_ of 44 nM, BZQ an IC_50_ of 1.05 µM and TCBZ, the only compound that killed the parasites in our assay, had the highest IC_50_ with 1.5 µM (Fig 5).

**Fig 5.**
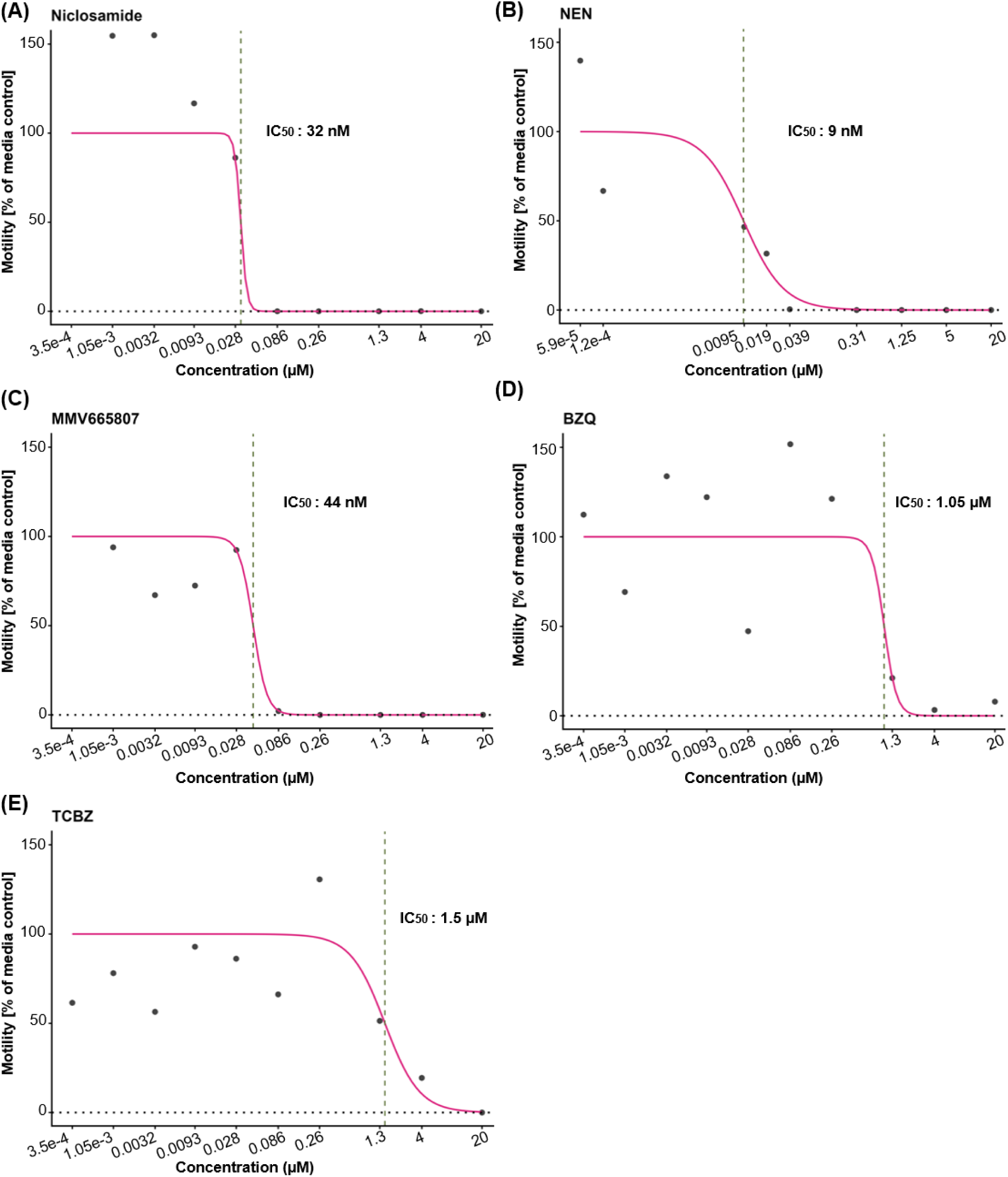
Dose-response curves of the most active compounds. NEJ motilities normalized with the media-only control are shown for increasing concentrations of niclosamide (A), NEN (B), MMV665807 (C), BZQ (D) and TCBZ (E). Data is derived from one biological replicate consisting of two technical replicates.

SEM images were collected of NEJ incubated for 72 hours with the active compounds and showed no morphological alterations at the concentrations corresponding to the IC50 with a phenotype closely resembling the DMSO control (Fig 6). In contrast, parasites exposed to 20 µM of niclosamide, NEN and MMV665807 appeared collapsed with a wrinkled surface (Fig 6A, B and C). Interestingly, while still alive and in culture, NEJ treated with niclosamide, NEN and MMV665807 appeared swollen and enlarged when visualized by light microscopy. NEJ exposed to 20 µM of BZQ lost their elongated shape and appeared rounded in SEM (Fig 6D), which reflects the phenotype observed by light microscopy in the treated live parasites.

**Fig 6.**
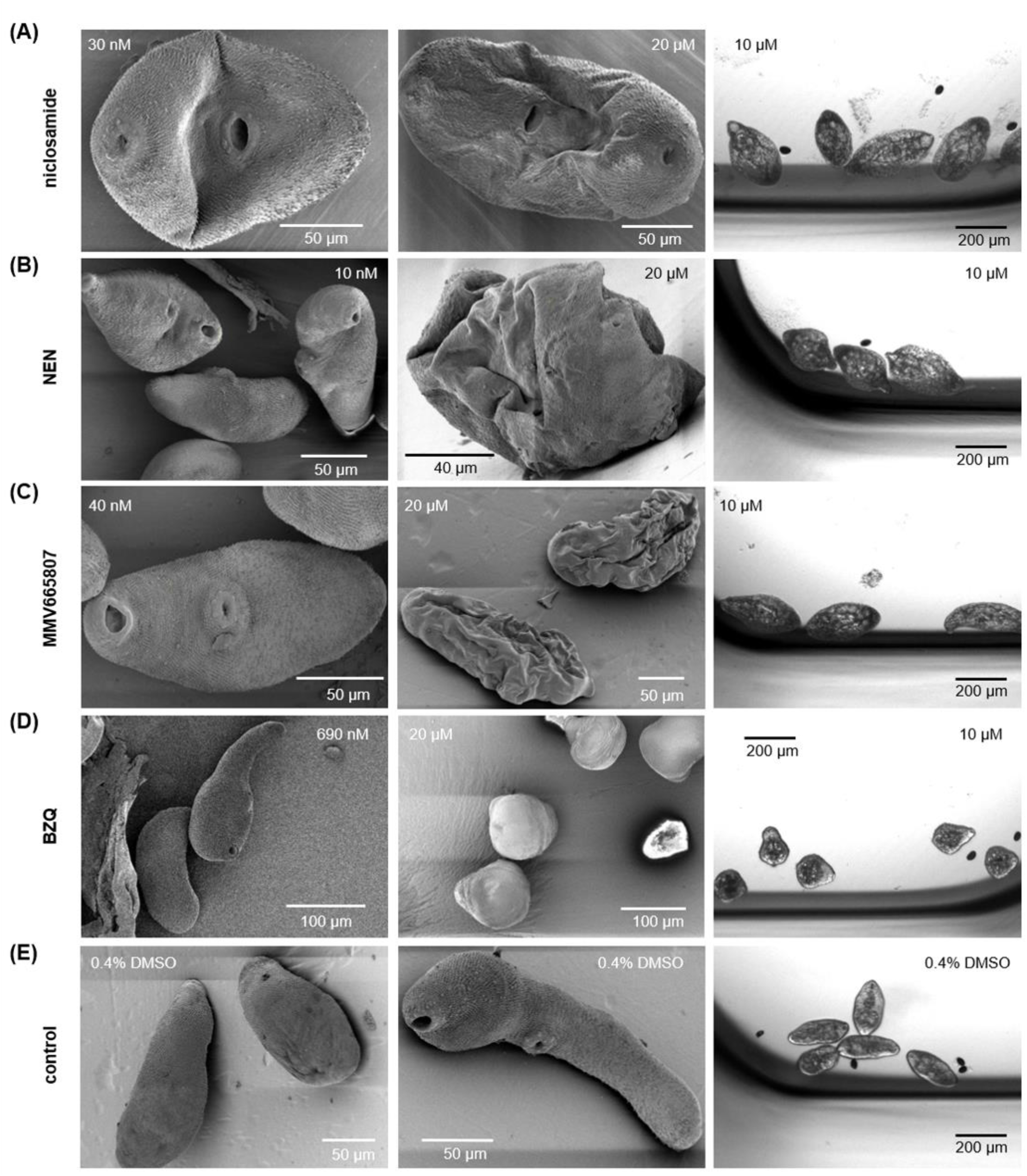
SEM and light microscopy images of drug treated NEJ. Parasites treated with niclosamide (A), NEN (B), MMV665807 (C), and BZQ (D) were imaged by SEM after exposure to two different drug concentrations and by light microscopy after exposure to 10 µM. Control parasites (E) were exposed to 0.4% DMSO. Drug treatments were carried out for 72 hours before imaging.

## Discussion

The discovery of new treatment options for fasciolosis has become a priority due to the limited number of available drugs and the emergence of resistant *F. hepatica* isolates. We propose a simple approach to access compound activity on *F. hepatica* NEJ based on quantification of parasite motility. The five-parasite-per-well plate setup enabled reliable identification of active hits, despite some variability in motility scoring. The relatively quick imaging with four frames per well every 10 seconds allows for increased throughput as compared to visual read-outs, and for objective quantification of parasite movement. Assay setup still requires manual pipetting of parasites which might be the main factor limiting throughput. The 72 hours incubation was chosen to identify compounds with a slower onset of action; this duration does not negatively affect overall parasite motility as determined in the media and DMSO controls. While the TCBZ assays served to establish the right assay conditions, they also revealed a time- and dose dependent activity profile for the reference compound TCBZ.

Amongst the panel of assessed anthelmintics, the onset of action at 10 and at 40 µM was slower for buparvaquone and BZQ compared to TCBZ and the salicylanilides (niclosamide, NEN and MMV665807). Buparvaquone is used in veterinary medicine against the protozoan parasite *Theileria annulata* [40] where it targets the Q_0_ site of mitochondrial cytochrome b [41], it was further shown to be active on the helminth *E. multilocularis* and to inhibit its mitochondrial respiration [36]. We did not prioritize buparvaquone due to its slow and incomplete activity in *F. hepatica*. BZQ showed a strong activity after 48 hours and brought parasite movement close to zero. It binds to an ubiquitously conserved residue in the praziquantel target TRPM ion channel and activates it, leading to increased Ca^2+^ signal and spastic paralysis of the parasites and tegumental disruption in juvenile and adult *F. hepatica* [34]. Our observations of contracted NEJ after BZQ treatment (light microscopy and SEM) are consistent with its known mode of action. As anticipated, praziquantel showed no activity against *F. hepatica*. The drug is effective against many platyhelminthes and the only treatment for the blood fluke *Schistosoma* [42]; however, a single amino acid substitution in the TRPM ion channel of *F. hepatica* renders the parasite insensitive to its effects [34]. Albendazole, mebendazole and fenbendazole are part of the benzimidazole class and are commonly used in veterinary and human medicine for their large spectrum activity against several helminths species [43,44]. Albendazole is used to treat the more mature stages of *F. hepatica*, however, none of these three benzimidazoles showed a clear effect on *F. hepatica* NEJ movement in our screen. Niclosamide, NEN and MMV665807 all belong to the halogenated salicylanilide class of pharmacologic molecules, such as the flukicides closantel, oxyclozanide and rafoxanide used to treat fasciolosis in livestock [45]. These compounds contain a β-hydroxy-carbonyl motif that has the potential to bind to multiple biological targets leading to pleotropic biological activities [46]. The best-studied amongst them is uncoupling of the energy generative oxidative phosphorylation [47,48]. Previous work done on tegumental NA^+^/K^+^-ATPase activity in *Fasciola hepatica* [49] highlighted the importance of these ion pumps in the osmoregulation of the parasite. The distortion of energy generative oxidative phosphorylation by salicylanilides could decrease the ATP production and affect the ion pumps at the surface of the tegument, leading to an influx of water in the parasite. During microscopic imaging of NEJ exposed to niclosamide, NEN and MMV665807, we observed pronounced swelling consistent with the proposed mode of action of these compounds, in which aberrant osmotic influx of water leads to parasite distension. Subsequent preparation for SEM resulted in dehydration of the previously swollen specimens, producing the characteristic collapsed and wrinkled morphology observed in the SEM images. This is in agreement with previous work on *Haplorchis taichui*, an intestinal trematode, that showed swelling on the tegumental surface after niclosamide treatment [50]. Halogenated salicylanilides currently used to treat fasciolosis primarily target more mature parasite stages; interestingly, the salicylanilides tested in this study exhibited strong activity against NEJs, with very low IC_50_ values.

Despite the pronounced inhibition of parasite movement and morphological alterations after drug treatment, niclosamide, NEN, MMV665807 and BZQ did not kill the parasites within 72 hours of incubation even at high concentrations. It remains to be investigated whether (1) parasites can recover after drug treatment and (2) if (temporary) inhibition of NEJ movement is sufficient for *in vivo* efficacy. Taken together, this work provides a reliable method to screen compounds for activity on *F. hepatica* NEJ and reveals three salicylanilides with previously unrecognized potent activity against the parasite.

## Acknowledgements

We thank Regula Flückiger for technical support. We are grateful to the Microscopy Imaging Center of the University of Bern (MIC), Bern, Switzerland for providing the microscopic infrastructure for parasite live imaging. We thank Daniel Sprague & Jonathan Marchant from the Department of Cell Biology, Neurobiology, and Anatomy, Medical College of Wisconsin, Milwaukee, WI, USA, for providing BZQ. We thank Verena Elbert and Gabriela Knubben-Schweizer from the Ruminant Clinic of the Ludwig-Maximilians-University, Munich, Germany, for providing *F. hepatica* metacercariae for pilot tests. We are grateful to the staff of the abattoir Bell in Oensingen, Switzerland for providing fresh infected livers. We further thank the Swiss National Science Foundation that provided the funds for this work (grant number 216088 awarded to NW) and the salary of MP (grant number 320030-227438 awarded to BLS). We are grateful to CIBERINFEC – CIBER de Enfermedades Infecciosas (CB21/13/00056), Instituto de Salud Carlos III (ISCIII), Ministry of Science and Innovation, and the European Union, for funding the research visit of GC under the program “Ayudas a la Formación y Movilidad”. The funders had no role in study design, data collection and analysis, decision to publish, or preparation of the manuscript.

## Supporting information

**Supplementary Video S1. Parasite movement over time.** Parasite movement of one representative well with five parasites is shown for four minutes. The left-hand side shows the microscopic images of the parasites, and the right-hand side shows the subtraction of pixel intensities between subsequent images as carried out to quantify parasite motility.

## Supporting information

**Fig S1.**
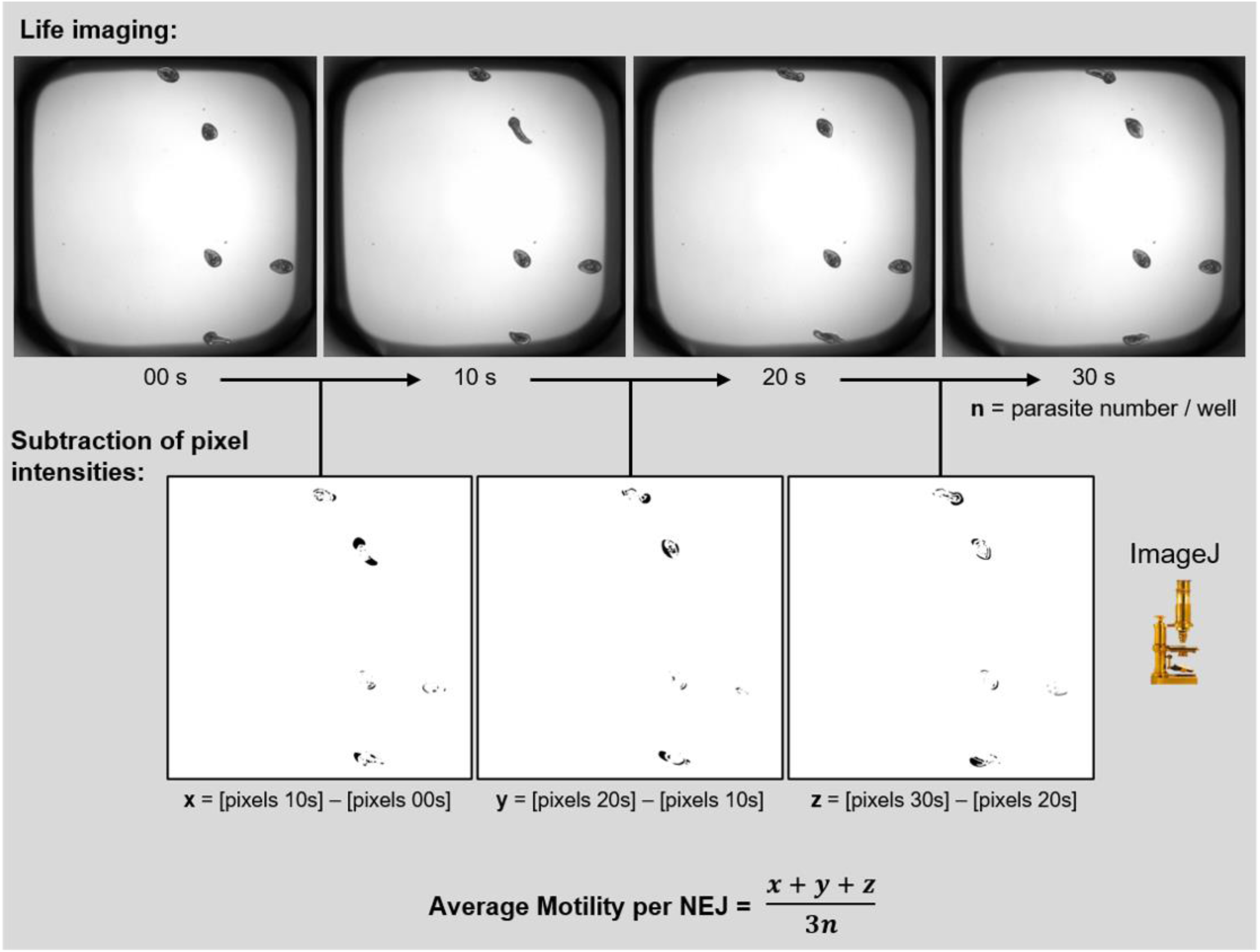
Quantification of NEJ motility with live imaging. Live imaging, image analysis, and calculation of the parasite motility are shown.

**Fig S2.**
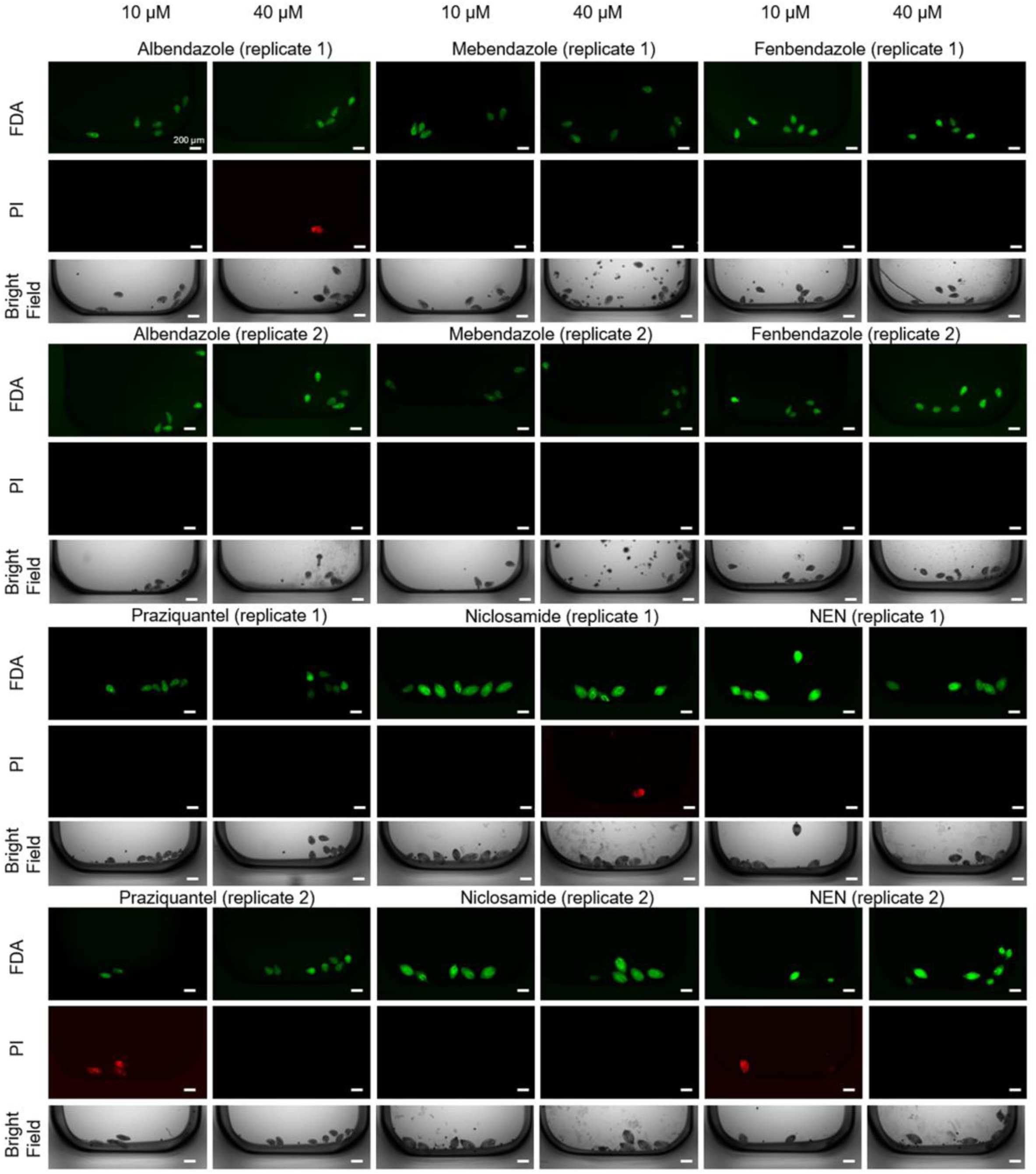
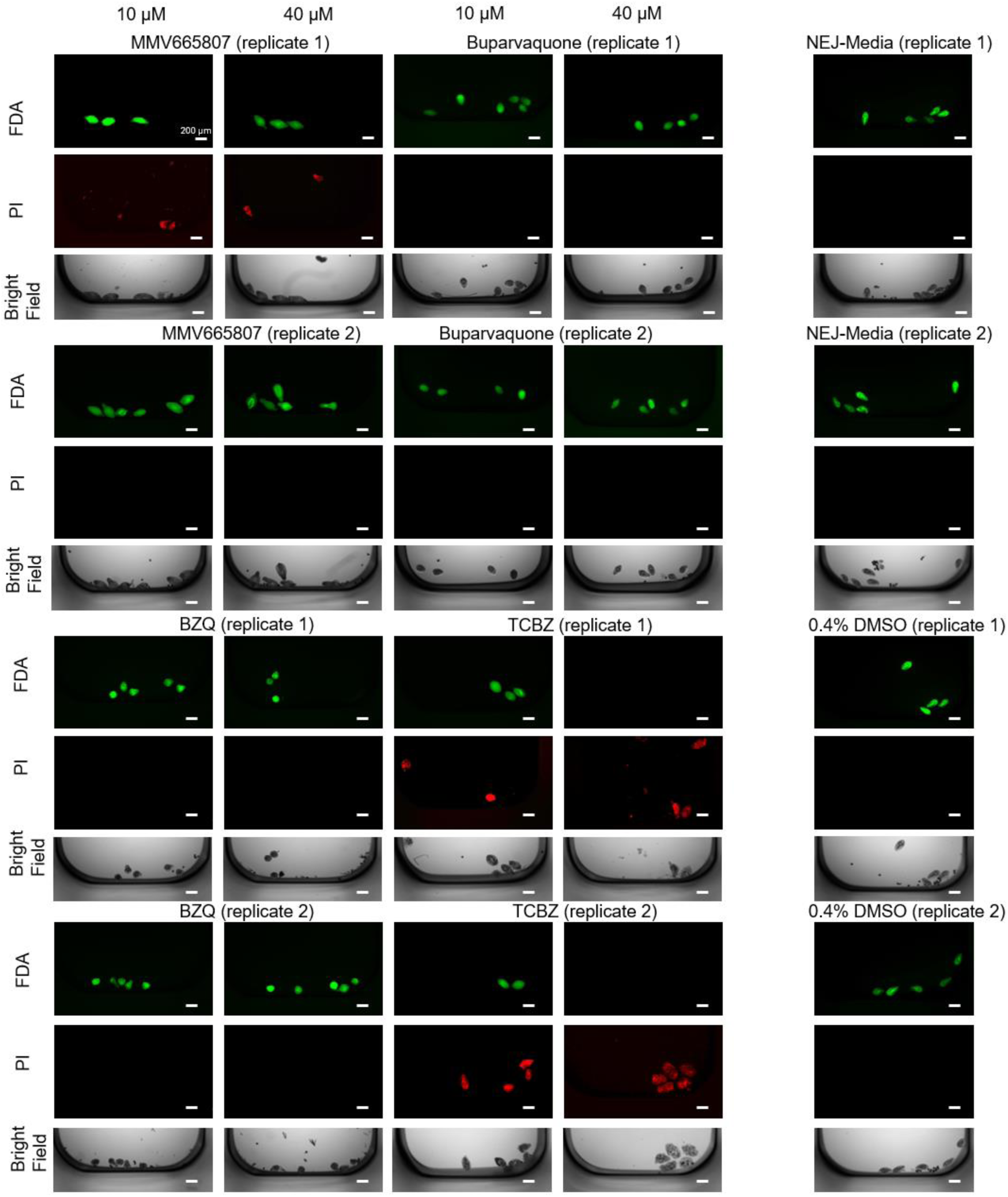
NEJ viability after drug exposure. Live/dead staining was performed after 72 hours of incubation with albendazole, mebendazole, fenbendazole, praziquantel, niclosamide, NEN, MMV665807, buparvaquone, BZQ and TCBZ, each applied at 10 and 40 µM. Data is derived from one biological replicate (one excystation) with two technical replicates (wells) per drug and concentration. Scale bars correspond to 200 µm.

